# Identification of universal and cell type specific p53 DNA binding

**DOI:** 10.1101/177667

**Authors:** Antonina Hafner, Lyubov Kublo, Galit Lahav, Jacob Stewart-Ornstein

**Affiliations:** Department of Systems Biology, Harvard Medical School, Boston, MA 02115, USA; Department of Computational and Systems Biology, University of Pittsburgh Medical School, Pittsburgh PA, 15260; Department of Developmental Biology, Stanford University, Stanford, CA 94305, USA

**Keywords:** p53, ChIPseq, Chromatin, DNA damage, Gene Expression

## Abstract

The tumor suppressor p53 is a major regulator of the DNA damage response and has been suggested to selectively bind and activate cell type specific gene expression programs, however recent studies and meta-analyses of genomic data propose largely uniform, and condition independent, p53 binding. To systematically assess the cell type specificity of p53, we measured its association with DNA in 12 p53 wild-type cell lines, from a range of epithelial linages, in response to ionizing radiation. We found that the majority of bound sites were occupied across all cell lines, however we also identified a subset of binding sites that were specific to one or a few cell lines. Unlike the shared p53-bound genome, which was not dependent on chromatin accessibility, the association of p53 with these atypical binding sites was well explained by chromatin accessibility and could be modulated by forcing cell state changes such as the epithelial-to-mesenchymal transition. These results position p53 as having both universal and cell type specific regulatory programs that have different regulators and dependence on chromatin state.

## Introduction

p53 is the major transcription factor regulating the DNA damage response (DDR) in mammals, by inducing transcription of genes involved in DNA repair, cell cycle arrest and apoptosis (1,2). Though ubiquitously expressed across human tissues, it remains unclear to what extent p53 functions are shared across different cell types. Context specific regulation of gene expression by p53 has been a long-standing hypothesis in the p53 field, and implies that p53 can integrate information about cellular context and the type of stress to selectively activate some target genes versus others (1,3–5). However, studies querying p53 binding genome wide in different human cell lines and upon different treatments have variously found strong agreement in p53 binding locations (6,7) and activation of a core set of target genes (8). Other studies have argued for unique cell type or transformation status specific p53 activity (9–11). These studies compared pairs of cell lines or supplemented single cell line data with meta-analysis of published datasets, an approach that is powerful for identifying universal p53 binding sites but has limits for detection of cell line specific binding patterns due to divergent experimental conditions across datasets.

In this work, we explored cell type and stimulus specificity of the tumor suppressing transcription factor p53 at the level of DNA binding. To study how p53 binding varies across cell lines, we measured p53 DNA binding in 12 cell lines from different tissue types in response to a single treatment. Indeed, we took advantage of our previous characterization of these cell lines which showed a comparable acute response of p53 (12) in response to ionizing radiation (IR). By treating this panel of epithelial cell lines with a dose of IR sufficient to induce uniform p53 activation across cell lines and measuring p53 binding at an early (2 hours) time-point we minimized secondary effects and focused on measuring the rapid and direct binding of p53. Our approach differs from the majority of p53 datasets in the literature, which use chemotherapy agents such as doxorubicin or the p53 activator Nutlin3A at later time-points of 6-12 hours. This coherent set of samples allowed us to rigorously explore the heterogeneity of p53 binding and identify the influence of universal genomic and cell line specific chromatin factors on p53 binding.

We found that the majority of p53 binding events to be universal across cancerous and non-cancerous cell lines, with strong quantitative agreement in binding magnitude. We further found that Nutlin3A treatment resulted in a nearly identical set of p53 binding events as IR, suggesting the conservation of these binding sites across treatments(6). The presence of highly conserved p53 DNA binding site is consistent with previous meta-analysis of p53 DNA binding (6). However, we also identified a set of variable p53 binding events (~5%) present in only one or a handful of cell lines. These binding events were often near transcriptionally active genes and correlated strongly with cell line specific chromatin accessibility. Consistent with this, we were able to alter p53 DNA binding when we pharmacologically modified the chromatin state or induced an epithelial-to-mesenchymal transition to globally change cell state. Taken together, our data shows that the majority of p53 DNA binding is context independent but there is a small but potentially important set of cell type or cell state specific binding sites for p53.

## Methods

### General genomic analysis

All DNA reads were single end Illumina reads and were aligned to HG19 genome build using bowtie (13). Aligned reads were further processed using HOMER (V4.6, (14)) to assemble tag files and call peaks. Peak locations and tag files were read and integrated with other datasets/types by a custom Matlab code (Mathworks). For each ChIP-seq dataset the number of reads in p53 peaks were normalized to the average of all cell lines, and for subsequent analyses and comparisons, peaks with less than 2 normalized counts were discarded. Clustering and comparisons were based on Pearson correlations between p53 binding signals. RNA data was aligned to the Refseq HG19 transcriptome using Tophat, CuffQuant, and CuffMerg (15) or Salmon (16). Genomic binding and signals were visualized using the UCSC genome browser (17). Motif analysis was performed in Matlab on the HG19 genome using a ChIP-seq derived PWM adjusted to have a minimum probability of occurrence for each nucleotide.

### ATAC-seq

ATAC-seq was performed as described (18), with the major exception of the use of a MuA transposase (Thermo) rather than the TN5 transposase. Briefly, MCF7 or LOXIMVI cells were trypsinized and 50K cells, spun down, washed once with PBS, and lysed with a hypotonic buffer containing 0.1% NP-40, and spun down to generate a crude nuclei pellet. This pellet was transposed in a 30μl volume using MuA (0.7μl), MuA buffer (10μl), and H2O (19μl) for 5min at 30C. The sample was treated with 3μl stop solution, and incubated at 30C for a further minute. The sample was then collected and purified by addition of 45μl of SPRI beads (Aline Biosciences). The purified sample was PCR amplified in two steps to add barcoded adaptors suitable for Illumina sequencing. Samples were sequenced with single end 75bp reads on an Illumina NextSeq. Reads (>30M) were trimmed to remove adaptors with cutadapt (19), aligned to the genome with Bowtie, and analyzed with Matlab. Genomic DNA (50ng) from MCF7 and LOXIMVI was transposed, amplified and sequenced in parallel to estimate background.

### ChIP-seq

P53 ChIP-seq was performed largely as previously described (20), briefly, 10M cells were treated with 4Gy IR (RS-2000, RadSource) and 2 hours later were fixed by addition of 1% paraformeldehyde (Alfa Aesar) at room temperature for 10 minutes with agitation. Fixation was stopped by addition of 250mM glycine. Cells were scraped and flash frozen. Cell pellets were thawed in hypotonic lysis buffer and spun to generate a crude nuclei prep. These nuclei were lysed in an SDS buffer and sonicated (Bioruptor) to fragment DNA. Fragmented DNA was diluted in IP buffer and agitated overnight with 2mg/ml DO-1 (anti-p53, Santa Cruz). 20μl of protein A magnetic beads (Invitrogen) were used to isolate the p53 associated fragments and samples were washed with low salt, high salt, and LiCl buffers. DNA was eluted from beads with an SDS/NaCO3 buffer and was de-cross-linked at 65C for 6hrs in a high salt buffer.

For experiments in Fig. 5, ChIP-seq was preformed using a Micrococcal Nuclease protocol. Briefly, cells were fixed and nuclei extracted as above, DNA was fragmented by a 20 minute incubation with Micrococcal Nuclease (NEB) at 37C. Nuclei were then lysed by brief sonication (Branson) and fragmented DNA were Immuno-precipitated as described above.

ChIP libraries were constructed with the commercial NEBnext kit (NEB) and associated protocols, although reaction volumes were reduced by 4 fold and custom adaptors and barcodes were employed. Libraries were sequenced with single end 75bp reads on Illumina NextSeq.

Reads were aligned to the HG19 genome with Bowtie1.1 (13), and analyzed with HOMER (14), MACS2 (21) and custom Matlab scripts. Peak calling was done after pooling reads (5-15M per line, ~150M total) from ChIP-seq experiments in all cell lines. The final set of peaks represented the consensus of HOMER (default settings) and MACS2 (using the q<0.01 threshold) identified peaks, and was filtered to remove ENCODE black-list locations. The number of reads within each peak region was computed from HOMER tag files using custom Matlab scripts. Background regions around each peak were subtracted from peak scores to correct for high background regions. The HOMER package (14) was used for *de novo* motif discovery. WebLogo was used to generate the motif plot (22) in (Figs. 1B, 2C) for the top enriched motif. The top enriched motif (Fig. 1B) was then used to re-scan and score all peaks and background regions. Background regions were generated by selecting 500bp regions adjacent to either side of the peak and excluding regions that overlap with p53 peak regions. Clustering of peaks was accomplished using a correlation distance metric and average linkage in Matlab.

**Figure 1:**
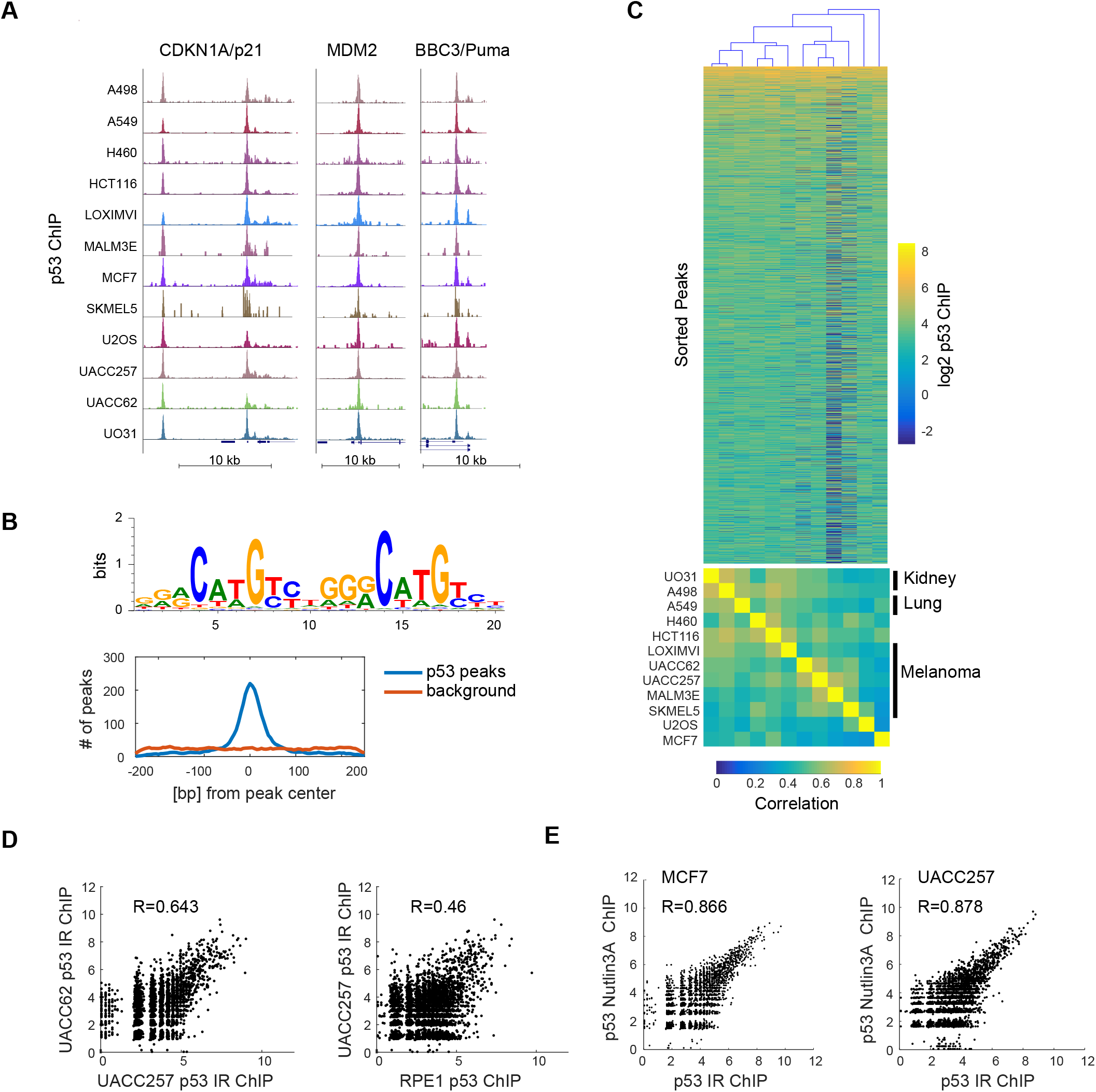
Stereotyped p53 binding across 12 cell lines. **(A)** ChIP-seq for p53 in 12 p53 wild-type cell lines. UCSC screen shots of p53 binding sites for three canonical p53 target genes are shown. **(B)** Motif analysis recovered a p53 motif that was centrally enriched within peaks. **(C)** Heatmap showing p53 binding intensity in 8742 locations in the genome. Cell lines were clustered on p53 binding and resulted in lineages clustering together. **(D)** Comparison of p53 binding in two cancer cell lines (UACC62 and UACC257) as well as between one cancer (UACC257) and one non-cancerous cell line (RPE1). **(E)** Comparison of p53 binding between Nutlin3A and IR treated samples in MCF7 or UACC257 cells.

**Figure 2:**
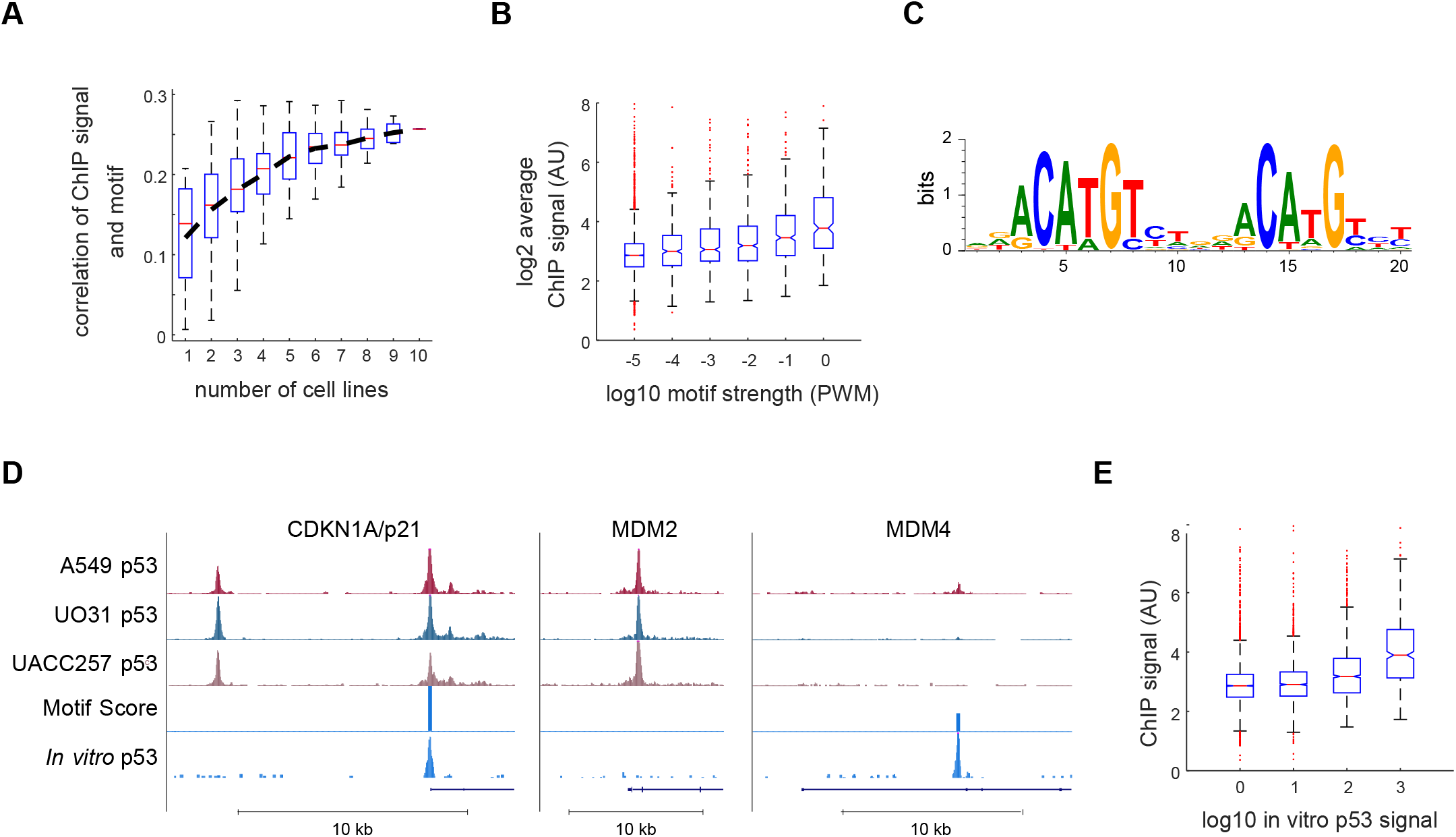
Genomic sequence is weakly predictive of p53 binding. **(A)** The correlation between motif strength and p53 binding is shown as a function of the number of cell lines across which the peak height was averaged, box plots represent the distribution of correlations across all possible cell line combinations. **(B)** The degree to which p53 binding motif predicts the strength of p53 binding is shown in a box plot, with p53 binding sites binned by their motif strength. **(C)** The top enriched motif identified by *in vitro* ChIP. **(D)** UCSC screenshots of p53 binding sites in A549, UO31 and UACC257 in response to IR, motif score, and *in vitro* p53 binding signal are shown for CDKN1A/p21, MDM2, and MDM4. **(E)** *In vivo* p53 binding strength is shown in a box plot, binned by *in vitro* p53 binding signal at each genomic site.

### *In vitro* ChIP-seq

To generate recombinant p53 we in vitro transcribed/translated human p53 with a c-terminal HA tag using a rabbit reticulocyte system (Promega). To generate fragmented genomic DNA we tagmented 50ng of human genomic DNA from MCF7 cells using the MuSeq kit (Thermo) and amplified it using PCR and custom adaptor primers for 8 cycles. DNA was cleaned up on SPRI beads (Aline Biosciences) and quantified. At room temperature 20ng of DNA and recombinant p53 (0.1uM final) were combined in a binding buffer (10mM TRIS, 5mM MgCl2, 10% glycerol, 1mM DTT) and incubated at room temperature for 30 min. The mixture was diluted 2 fold (to 20ul) and 1.5ul of anti-HA antibody was added (Rockland) and the sample incubated at 4C overnight with shaking. A 1:1 mixture of magnetic proteinA/proteinG beads was added (Sigma) and incubated at 4C for 1hr with shaking. The beads were then washed 3x with washing buffer (10mM Tris, 5mM HCL, 0.1% triton, 150mM NaCl) and DNA eluted with elution buffer (1%SDS, 100mM Na2CO3) at 37C for 15 minutes. Samples were cleaned up, and adaptors and barcodes added by PCR. Reads (>30M) were trimmed to remove adaptors with cutadapt (19), aligned to the genome with Bowtie, and analyzed with Matlab.

### RNA-seq

For each cell line 50,000 cells were plated in 35 mm dishes, 24hrs later cells were treated (or not) with 4Gy IR (RS-2000, RadSource), 3hrs after that cells were lysed with Trizol (Ambion). RNA was purified on affinity columns and DNAse treated (Zymo). Purified RNA (500ng) was polyA purified using magnetic beads (NEB), fragmented and reverse transcribed using protoscript RT (NEB), second strand synthesized (NEB), and then assembled into libraries with the commercial NEBnext kit (NEB) and associated protocols, although reaction volumes were reduced by 4 fold and custom adaptors and barcodes were employed. Libraries were sequenced with single end 75bp reads on a NextSeq.

### Cell Culture and Cell Treatment

Parental cell lines were obtained from ATTC, with the exception of the RPE cells which were a gift from Prof. Steve Elledge (Harvard Medical School), and were thawed and propagated in RPMI (GIBCO) with 5% FBS. All experiments were performed in this media. All media was supplemented with 1% antibiotic and antimycotic (Corning). Treatment with Nutlin3A (Sigma) was at 5uM. X-ray induced DNA damage was generated with a RS-2000 source (RadSource, 160KeV). MCF7 cells were treated with 2uM decitabine (5-AZA-2’-deoxycytidine, MP Biomedicals) for 5 days, cells were split on day 2, re-plated in decitabine containing media. Treated and untreated cells were then further treated with IR or not as with other samples. A549 cells were induced to undergo epithelial-to-mesenchymal transition by treatment with TGFB (Sigma) at 2.5ng/ml for 5 days. For knockdown of p53, A549 cells were infected with a doxycycline inducible p53sh (23), selected on puromycin for infected cells. Subsequent induction of doxycycline was for 24hrs with 500ng/ml (sigma).

### Public datasets

Raw fastq files were downloaded from the Sequence Read Archive (see supplement). These datasets were all single end Illumina reads and were aligned to the HG19 genome with using the same pipeline as described above for our ChIP-seq samples, and further analyzed with HOMER to generate tag files. Custom Matlab code was used to compare these datasets to our ChIP-seq data.

### Statistics

Statistics relating to motif enrichment or GO-term enrichment were from multiple hypothesis corrected hypergeometric tests performed by HOMER (for motif calling) or using Matlab. Correlation coefficients are Pearson unless otherwise noted and were assigned p-values by MATLAB using a two tailed t-test as sample sizes were sufficiently large (1000s).

## Results

### p53 binding across the genome is stereotyped between cell lines and treatments

To study how p53 binding varies across cell lines we treated 12 cell lines expressing wild type p53 with ionizing radiation (IR; X-Ray 4Gy) to induce p53 activity and 2 hours later cross-linked and harvested each cell line for ChIP-seq analysis. Visual inspection of well-established p53 target genes showed clear ChIP peaks in all 12 cell lines (Fig. 1A). Overall, by pooling data from all cell lines we confidently called ~9000 p53 ChIP peaks. *De novo* motif analysis identified the p53 binding motif as the strongest single motif and also found it to be centrally enriched within peaks (Fig. 1B).

The quantitative strength of p53 binding at each genomic locus was highly conserved across the 12 cell lines (Fig. 1C). Though no strong groups of cell lines appeared by eye, hierarchical clustering correctly sorted the cell lines by tissue of origin, with pairs of lung and kidney lines, and all five melanoma lines clustering together (Fig. 1C). These p53 bound regions were also similar to other published datasets (6)(average within dataset correlation 0.53, average correlation to external datasets 0.49; Fig. S1). It was previously suggested that cancer cell lines show a different p53 binding profile from non-cancerous cells (9). We therefore compared the 12 cancer cell lines to an identically treated non-transformed line, RPE1. We found that p53 binding at identified sites in RPE1 cells in response to IR was highly correlated with p53 binding in the 12 transformed cell lines (Fig. 1D; average R= 0.48 for correlation (RPE, Cancer Lines) vs 0.53 for correlation (Cancer, Cancer)).

To further explore if the apparent uniformity of p53 binding is specific to IR, we treated two cell lines, MCF7 and UACC257, with a small molecule, Nutlin3A, that is known to activate p53 (24). Comparison of ChIP peaks between different conditions and cell lines, showed that IR-Nutlin3A correlations within each line that were stronger than any line - line correlations (Fig. 1E; R=0.87 or 0.88 for MCF7 and UACC257, respectively, vs R=0.73 for the maximum line-line). Thus, in addition to the uniformity we observe between cell lines, IR induced and pharmacologically induced p53 do not lead to distinct p53 function as measured by acute p53 DNA binding.

### Genomic DNA sequence has limited predictive power for p53 binding strength

Given the strong conservation of p53 binding across cell lines, and the recent analyses showing that DNA sequence is the best predictor of genomic p53 binding (6) we wondered if the DNA sequence was predictive of p53 binding strength. We tested this by comparing motif scores (calculated from the position weight matrix (PWM)) with p53 ChIP-seq signal intensity. The extent of the correlation between p53 ChIP signal and PWM score was highly cell line dependent (Fig. 2A), ranging from no correlation to correlation of 0.22 in a single cell line. Averaging p53 binding over increasing numbers of cell lines resulted in better agreement between genomic motif score and p53 binding, with the highest correlation being 0.26, when we averaged across all datasets (Fig. 2 A, B). Therefore, although the motif score significantly correlates with p53 DNA binding (p=1.98e-132), it only accounts for ~6% of the variance.

To explore if our motif analysis was simply a poor model of p53 binding, we performed an *in vitro* ChIP experiment. In this experiment, recombinant p53 was incubated with fragmented genomic DNA. This was followed by immunoprecipitation and deep sequencing similarly to a recently published protocol (25). As this assay uses sheared genomic DNA (with a size of ~300-600bp), effects of chromatin or other factors that may influence *in vivo* p53 DNA interactions, should not be present. We obtained a strong signal of p53 binding which was reproducible between replicates (Fig. S2A, B), recovering a consensus p53 motif (HOMER p=1e-2422, Fig. 2C), very similar to the motif found *in vivo* (Fig. 1B). We observed p53 binding sites, such as the one proximal to the CDKN1A/p21 promoter, that showed strong *in vivo* binding, a strong motif, and substantial *in vitro* p53 binding (Fig. 2D). Surprisingly, other binding sites, such as the one contained in the first intron of MDM2, showed substantial *in vivo* binding, but little *in vitro* binding and no strong motif. Conversely, the binding site at the MDM4 gene showed strong *in vitro* binding and a strong motif, but little *in vivo* binding. Overall, the *in vitro* p53 binding signal did not show a better correlation (R=0.252, p= 3.10e-127, Fig. 2E) with *in vivo* p53 binding than the motif score although we note that in this case measurement noise of in vitro binding likely degraded this correlation. These results suggest that factors other than DNA sequence determine p53 binding *in vivo*.

### A subset of p53 binding sites are cell-type specific

Our finding of a uniform set of p53 bound regions independent of cell line or even treatment is consistent with previous work (6). However, we wondered if we could also find cell-type specific p53 binding that due to the uniformity of our dataset (both in treatment and data collection) and early time-point of treatment, might have been missed in earlier analyses. We compared the cell line to cell line variability in p53 ChIP signal after correcting for the average ChIP peak signal (which contributes shot noise to our analysis) and identified about 5% of peaks (494 peaks) that showed high variation between cell lines relative to their average peak strength (Fig. 3A, B). For example, p53 peaks nearby the inflammatory associated genes IL1A and CXCL1 showed clear p53 binding in the LOXIMVI line, weaker association in the UO31 and H460 lines, and no binding in other cell lines (Fig. 3B). We also found variability in p53 binding at the promoters of previously reported p53 target genes, ALDH3A1 and EPHA2, ranging from no binding in some cell lines to strong peaks in others (Fig. 3B). *De novo* motif search on this set of variable peaks identified the p53 binding site as the most significantly enriched motif (HOMER, p=1.0e-46), suggesting that these sites represent direct p53 binding events.

**Figure 3:**
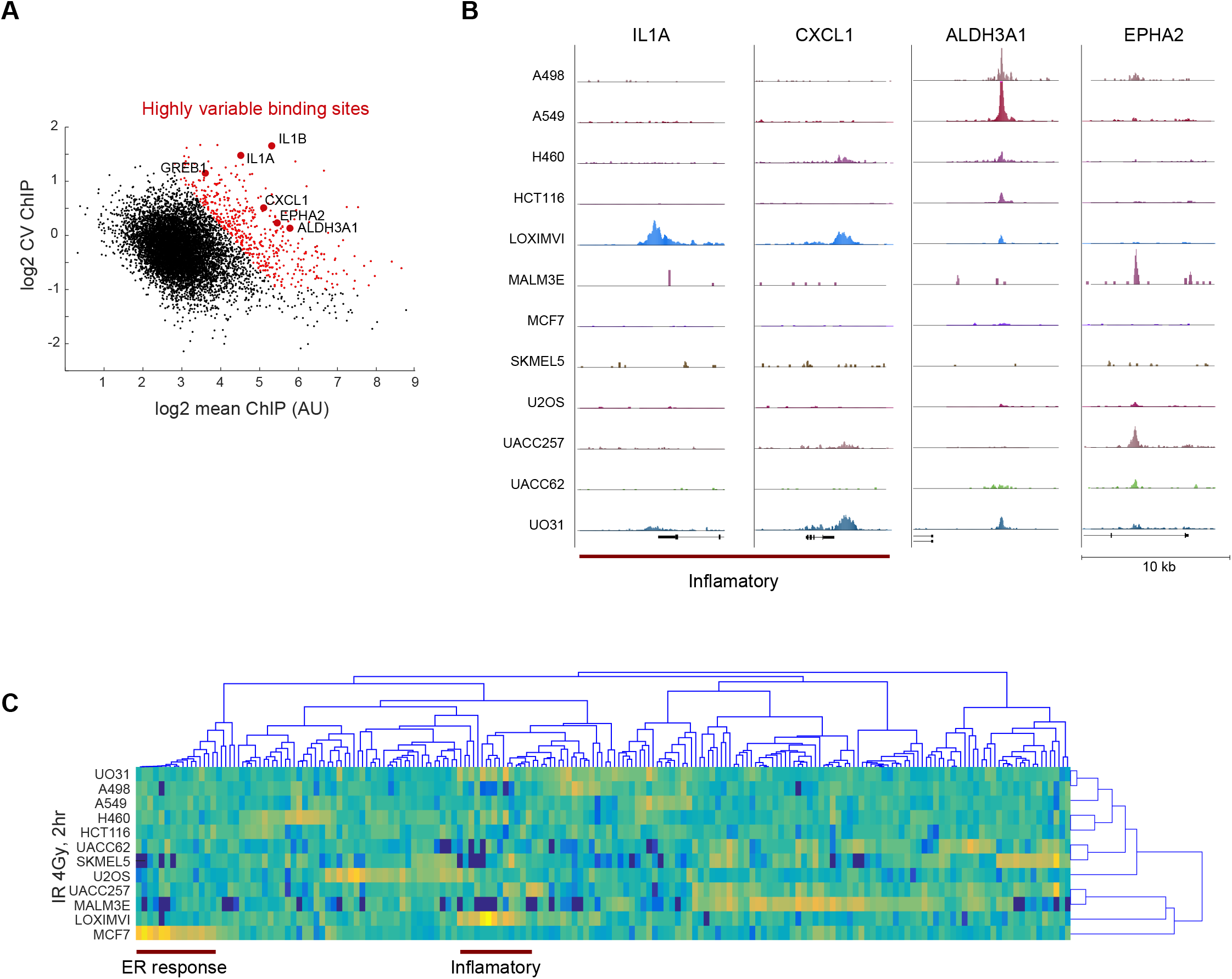
Variable p53 binding sites show cell type specific functional enrichment. **(A)** Scatterplot of all 8742 p53 binding sites by their average ChIP signal and coefficient of variation (CV). Highlighted in red are ‘highly variable’ peaks defined as having higher than expected CV relative to the peak height. Example binding sites are labeled with the associated gene names. **(B)** UCSC screen shots of four example ‘variable’ peaks. **(C)** Heatmap of ‘variable’ p53 peaks that are also nearby (<10kb) transcription start sites of genes. The intensity of each peak is normalized to the average across 12 cell lines. Cell lines and peaks were hierarchically clustered, with no grouping by lineage observed for cell lines. Groups of inflammatory and the ER associated are highlighted.

To determine if these highly variable binding sites had novel cell line specific functions, we assigned each peak to its closest gene (with a 10 kb cutoff) and clustered the resulting 218 peaks on their p53 binding. We found that most cell lines showed a few unique p53 binding peaks, but without strong clustering between cell lines (Fig. 3C) as in Fig. 1C. Enrichment analysis identified inflammatory/chemotaxis associated genes as being enriched in these highly variable p53 bound genes. The cell line LOXIMVI showed particularly strong enrichment for p53 binding to inflammatory genes including IL1A, IL1B, CLL20, and CXCL1 genes. UO31 also showed substantial binding for many of these targets. We also observed, that in the estrogen receptor (ER) positive MCF7 breast cancer cell line, several unique binding sites overlapped with ESR1 (estrogen receptor) binding sites, including TFF1, IGFBP4, and PRLH. These results suggest that the non-uniformities in p53 binding we observed may be linked to cell line specific regulatory programs.

### Cell line specific chromatin accessibility accounts for variability in p53 binding sites

The differences we observed between *in vivo* and *in vitro* DNA binding and the presence of cell type specific p53 binding, cannot be explained by DNA sequence. We thus hypothesized that chromatin accessibility may play a role in tuning *in vivo* p53 DNA binding. Consistent with this hypothesis, we observed a high correlation of cell-line specific p53 peaks with basal gene expression (p=1.9e-31), that we measured by RNA-seq in the 12 cell lines we studied. For example, expression of IL1A, IL1B, CXCL1, and GREB1 were all associated with p53 binding across the 12 cell lines (Fig. 4A). It has been suggested that p53 can act as a pioneer factor with a high affinity for histone occupied regions (26–28), whereas others have shown that p53 binds readily in open regions (29,30). Our results linking basal expression of nearby genes to p53 binding suggest that the ‘openness’ of the genomic region might influence p53 binding.

**Figure 4:**
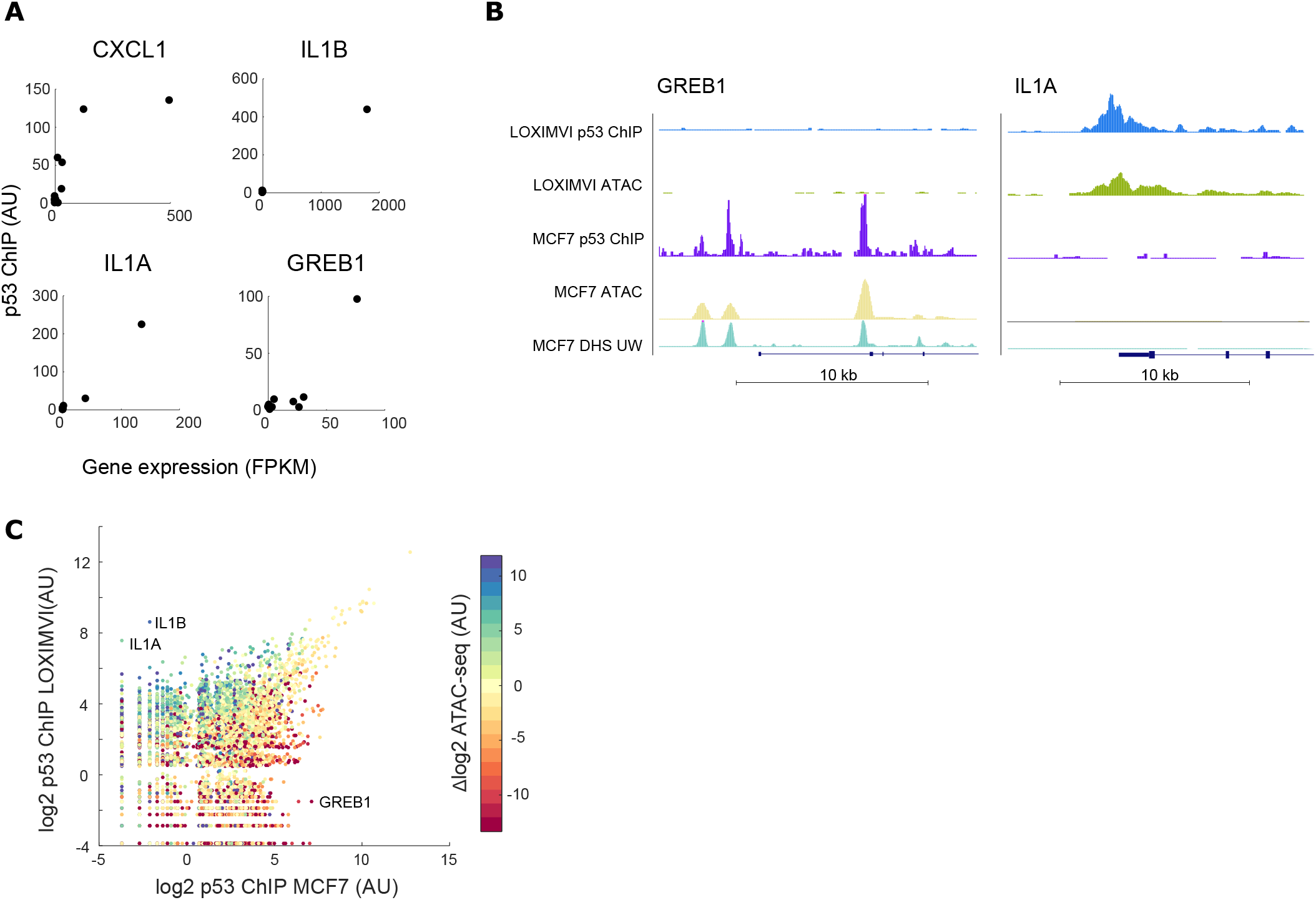
Chromatin accessibility contributes to variable p53 binding. **(A)** Scatterplots illustrating the relationship between basal gene expression and p53 binding across the 12 cell lines for four p53 peaks/genes (note that in many cases multiple cell lines show little gene expression or p53 binding and therefore cluster near the origin). **(B)** UCSC screen shots of two p53 binding sites; p53 binding in the proximity of GREB1 is found in MCF7 treated with IR, while IL1A was bound in IR treated LOXIMVI cells. ATAC-seq data and published DNase hypersensitivity data (for MCF7, untreated) showing that IR induced p53 binding correlates with basal DNA accessibility in each cell line. **(C)** Scatter plot of p53 binding post IR in MCF7 compared to LOXIMVI, colored by the difference in ATAC-seq signal: log2(LOXIMVI-MCF7) between the two cell lines.

To directly measure the connection between chromatin accessibility and DNA binding, we performed ATAC-seq. We chose two cell lines, LOXIMVI, which showed strong, and unique binding of p53 nearby inflammatory related genes and MCF7, which showed p53 binding at estrogen receptor associated genes. We performed a modified ATAC-seq protocol using the MuA transposase to generate genome wide maps of accessible regions in the MCF7 and LOXIMVI cell lines. Our ATAC-seq data and ENCODE produced DNAse sensitivity data from MCF7 showed strong overlap with greater than 90% of ATAC-seq peaks being Dnase accessible (31). We compared our ATAC-seq data to the p53 ChIP-seq signal for the inflammatory genes that showed p53 binding in LOXIMVI but not in MCF7 and observed strong ATAC-seq signal only in the LOXIMVI cell line (Fig. 4B), consistent with increased accessibility at these loci leading to stronger p53 binding. Conversely, GREB1, a breast cancer associated gene showed only p53 binding and ATAC-seq sensitivity in MCF7 cells (Fig. 4B). Moreover, genome wide, the difference in ATAC-seq signal between the two lines accounted for 22% of the variance in p53 binding between the two datasets (R^2^=0.225; Fig. 4C). More generally, as has been observed for other transcription factors (32), combining accessibility and motif scoring allows for improved prediction of DNA binding. Indeed, accessibility and motif score accounted for 13.8% and 20.9% of the variance in the log2(p53 ChIP-seq peak signal) for MCF7 and LOXIMVI respectively, compared to ~6% with the motif alone. We therefore conclude that chromatin accessibility favors p53 binding and accounts for a substantial fraction of the cell line specific gain of p53 DNA binding sites between MCF7 and LOXIMVI cells. Interestingly, we also found that genome wide chromatin accessibility was negatively correlated with *in vitro* p53 binding (R=-0.2, p=2.1e-80, MCF7 ATAC-seq vs. *in vitro* binding), suggesting that many strong p53 binding sites are obscured by local chromatin context.

### Perturbation of cell state alters p53 DNA binding

To establish a causal link between chromatin state and p53 binding, we treated MCF7 cells with decitabine, a methylase inhibitor that has been shown to broadly alter chromatin structure (33). We then treated these cells with IR and preformed p53 ChIP-seq and ATAC-seq on the samples. Comparing p53 binding between the decitabine treated and untreated cells, showed a modest but significant correlation between change in chromatin accessibility and change in p53 DNA binding between decitabine treated and untreated samples (R=0.16, p=3.99e-13). Looking at differential peaks between conditions, we found only one binding site, adjacent to the DLGAP5 gene, that showed a substantial change in p53 binding (Fig. 5A). This increase in p53 binding was accompanied by increased accessibility (Fig. 5B). The DLGAP5 binding site has a consensus p53 motif and showed occupancy in other cell lines such as UACC62 (Fig. 5B), consistent with this binding site being obscured by chromatin in wild type MCF7 cells. Overall, these data show that increased chromatin accessibility favors p53 binding but does not alter the global p53 DNA binding profile (Fig. 5A), perhaps due to limited overlap of accessibility changes induced by decitabine and p53 binding sites.

**Figure 5:**
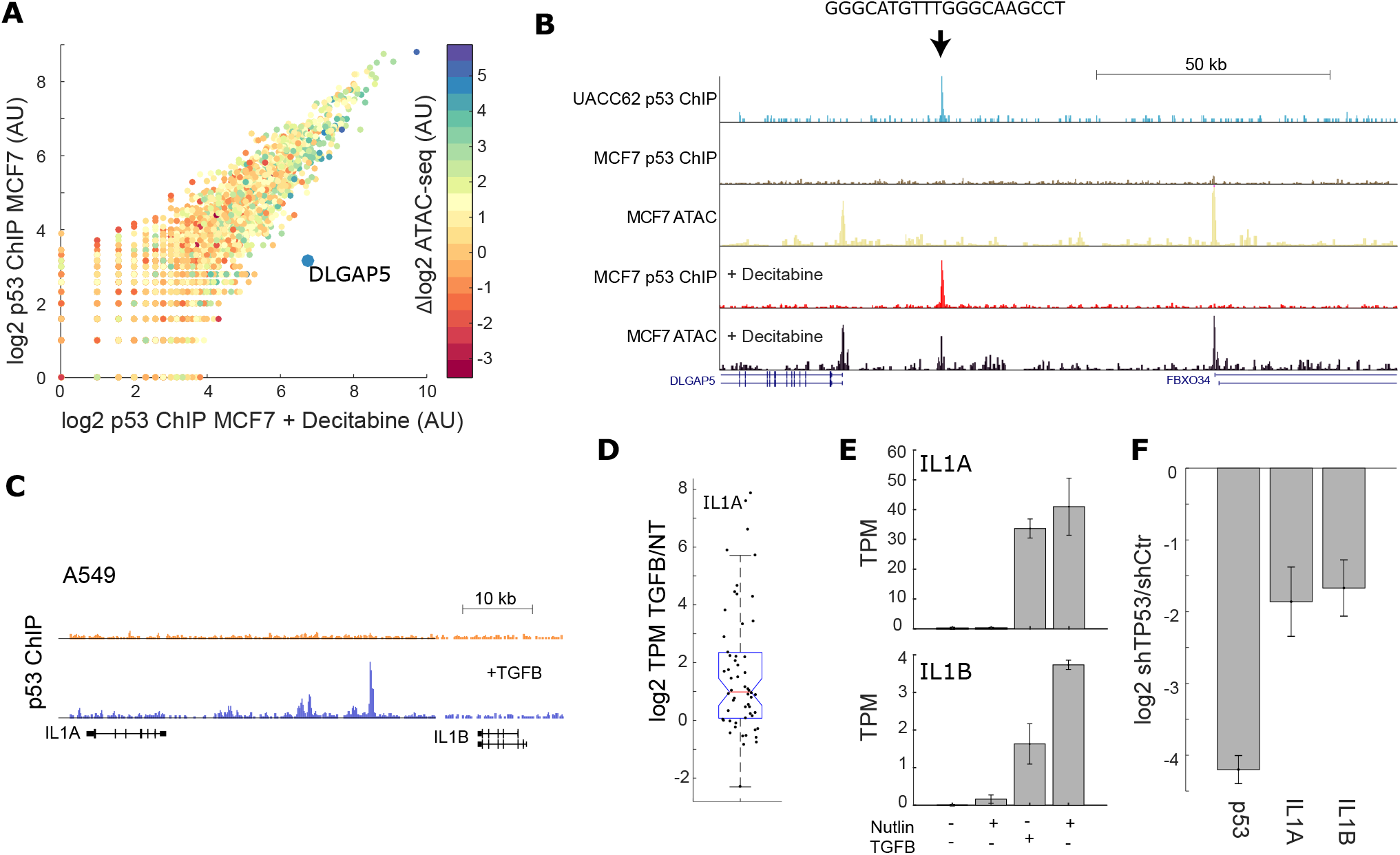
Cellular state regulates p53 binding. **(A)** Scatter plot of p53 binding in IR treated MCF7 cells compared to MCF7 treated with IR and decitabine, colored by the difference in ATAC-seq signal between decitabine treated and untreated cells: log2(decitabine-not treated). **(B)** UCSC screen shot of the region around the gene DLGAP5, showing changes in p53 binding and accessibility in the decitabine treated MCF7 cells (the new peak is indicated by an arrow). Binding of p53 in IR treated UACC62 cells at the DLGAP5 locus without decitabine treatment is also shown. **(C)** UCSC screen shot of the IL1 locus showing increased p53 binding in TGFB treated cells. **(D)** Boxplot showing log2 change in gene expression (TPM of TGFB treated/untreated) in genes nearby p53 binding sites that showed altered occupancy in TGFB treated cells. **(E)** Gene expression of IL1A and IL1B in cells treated as indicated with Nutlin3A or TGFB (N=3 experiments, except TGFB + Nutlin3A N=2). **(F)** Expression of IL1A and IL1B in A549 cells treated with TGFB comparing p53 knockdown cells to control knockdown (N=3). All error bars are SEM.

We next asked whether more dramatic perturbation of chromatin accessibility and cellular state can alter the cell type specific p53 DNA binding sites. The LOXIMVI cell line has gained p53 binding at inflammatory genes such as IL1, that we were unable to induce with decitabine treatment in MCF7 cells. We noted that the LOXIMVI line has been previous reported to have an EMT phenotype (34). We wondered if forcing an EMT transition on another cell line would result in changes to p53 binding. A549 cell have been shown to undergo EMT when treated with TGFB (35), we therefore treated A549 cells with TGFB for five days to induce EMT and measured p53 binding with ChIP-seq. We observed many new binding sites for p53, particularly we noted the emergence of p53 binding at the IL1A/B locus (Fig. 5C). Novel or stronger binding in the genome (2 std. dev. above untreated) was associated with increased expression of nearby genes under basal p53 condition (Fig. 5D). In particular, we noted increased expression of IL1A and IL1B under both basal p53 and Nutlin3A induced p53 (Fig. 5E). Expression of IL1A and IL1B was partially dependent on p53 as knockdown of p53 reduced expression of these genes (Fig. 5F), this was also true for the LOXIMVI line (Fig. S3) which endogenously expresses Il1A/B and has associated p53 binding at these genes.

## Discussion

The transcription factor p53 regulates the cellular response to DNA damage, including up regulating repair, cell cycle arrest and apoptotic proteins. The nature, strength, and balance between the DNA repair and cell death arms of p53 signaling varies across tissues in the body (3,12,36), and can be modified by drug treatment (37,38) and genetic perturbation (39). The role of the p53 itself in this decision making is controversial, with arguments for p53 behaving as a smart ‘signal integrator’ (reviewed in (1)) or a simple effector (6). We sought to understand the role of p53 in diverse cell lines by focusing on p53 DNA binding and gene expression in response to ionizing radiation.

As other studies have suggested, p53 DNA binding does not greatly vary across cell lines or treatments (6). However, we did find that p53 binding could group cell lines by their tissue of origin, suggesting some degree of tissue specificity. Further, we noted a modest, but significant correlation between the strength of p53 binding (measured by ChIP-seq) and the predicted strength of p53 association (using the PWM model). This correlation varied across cell lines and was strongest in the pooled dataset containing all cell lines. More strikingly, we observed a similar correlation when comparing genome wide *in vitro* association of p53 with *in vivo* p53 binding. In general, p53 binding at any given location in the genome was relatively poorly predicted by either *in vitro* binding or motif analysis suggesting that *in vivo* factors greatly contribute to p53 binding specificity.

Taking advantage of the coherence of our dataset we identified p53 binding sites that were variably occupied across cell lines. This subset of peaks were nearby genes enriched for specific cellular programs, most notably the inflammatory response in the melanoma LOXIMVI cell line and ER specific response in the MCF7 cell line. Our ATAC-seq data showed that this differential p53 binding could be attributed to differences in chromatin accessibility between the MCF7 and LOXIMVI cell lines. Globally, our data showed that a higher degree of chromatin accessibility favored p53 binding and is consistent with a previous report showing that open chromatin can provide a permissive environment for p53 (27) as well as other transcription factors such as ER (40) for example. However, as has been recently observed (41), p53 has strong ‘pioneer factor’ activity in some contexts; consistent with this we observe that many strong p53 binding sites such as those adjacent to CDKN1A show little chromatin accessibility prior to p53 activation.

We observe strong p53 binding to inflammatory genes in the LOXIMVI cell lines and also in the TGFB induced A549 line. Expression of these inflammatory genes was partially dependent on p53 (Fig. 5, Fig. S3). These results mirror an emerging role for p53 in inflammatory gene regulation in macrophages (42) and fibroblasts (43). Depending on the degree and context in which p53 drives these inflammatory signaling this may position p53 as a regulator of inflammatory signaling in epithelial systems including many cancers.

Taken together, these results suggest there may be two classes of p53 binding sites that are not clearly distinguished by p53 binding motif, those that require accessible chromatin or other auxiliary factors to function, and those where p53 is sufficient to bind and open chromatin. Supporting a mixed model of partial dependence of p53 on the cellular state to regulate its binding, we showed that alteration to the cellular state either using pharmacological agents targeting chromatin or the endogenous ligand TFGB to alter cellular state resulted in substantial changes to p53 binding. Further studies coupling chromatin accessibility, p53 binding, post-translational modifications, and measurements of RNA synthesis and degradation rates will be required to reconcile different models of p53 regulation and identify what features tune the cellular response to DNA damage in different cellular backgrounds.

## Data Access

All sequencing data has been deposited in NCBI’s Gene Expression Omnibus under accession number GSE100292. Data is also available as UCSC tracks as a custom session accessible at: https://genome.ucsc.edu/cgi-bin/hgTracks?hgS_doOtherUser=submit&hgS_otherUserName=jacobso&hgS_otherUserSessionName=Hafner_et_al_hg19_2019B

## Authors contributions

AH and JSO initiated and conceived the study; AH, LK, and JSO collected the data; AH and JSO analyzed the data; AH, GL and JSO wrote the paper, GL and JSO supervised the study.

## Acknowledgements

We thank Megha Padi and all members of the Lahav lab for feedback. We thank the Harvard Bauer sequencing core, Christian Daly, and Health Sciences Sequencing Core at Children’s Hospital of Pittsburgh for sequencing support and HMS research computing for maintaining the Orchestra/O2 cluster.

---

**Figure S1:**
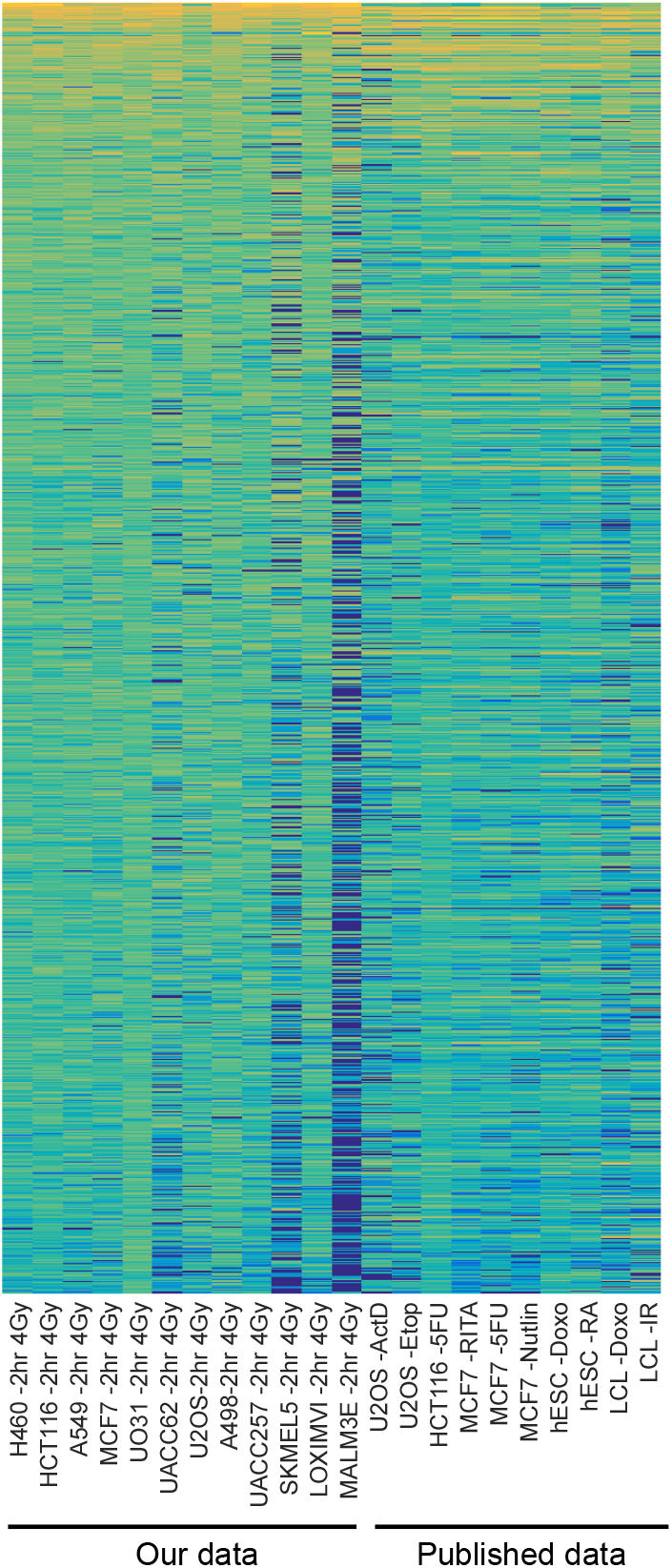
Comparison of p53 DNA binding with published p53 ChIP-Seq datasets. Heatmap showing p53 binding intensity in 8742 locations in the genome in 12 IR treated cell lines (same as Fig. 1C) as well as published datasets.

**Figure S2:**
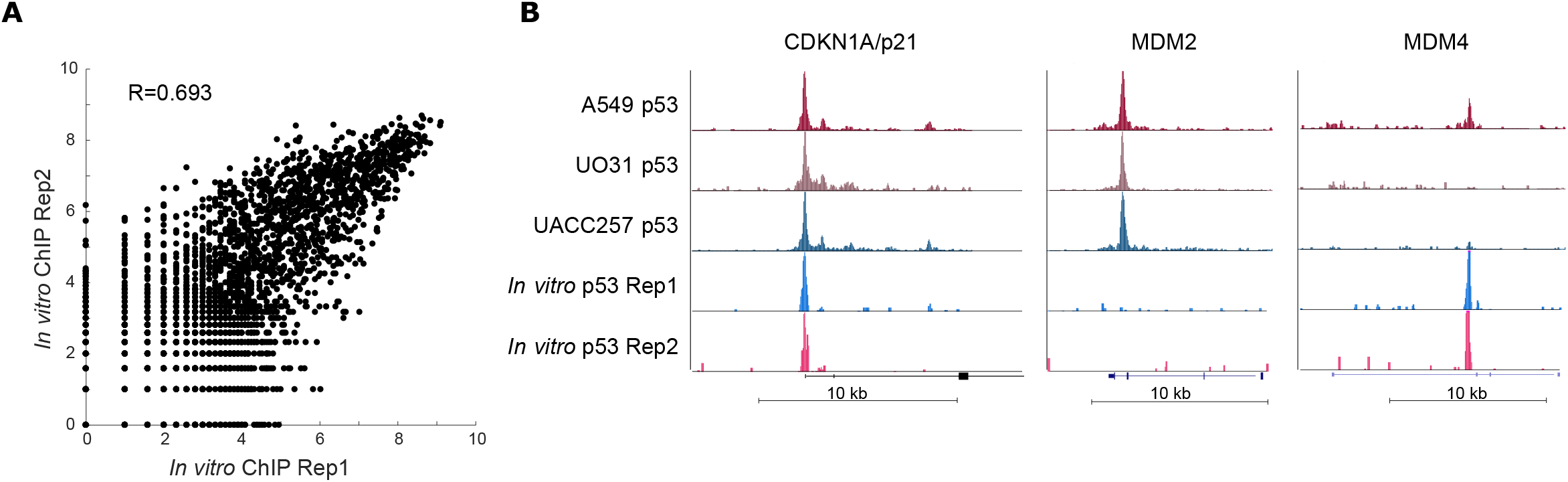
In vitro measurements of p53 are reproducible and show p53 sequence preference is partially recapitulated in vivo. **(A)** Quantitative agreement in binding strength at p53 binding sites between two replicate p53 *in vitro* IP datasets using different p53 protein preps. **(B)** UCSC browser shots of three key binding sites for p53 showing agreement between *in vitro* binding datasets and divergence of *in vivo* data.

**Figure S3:**
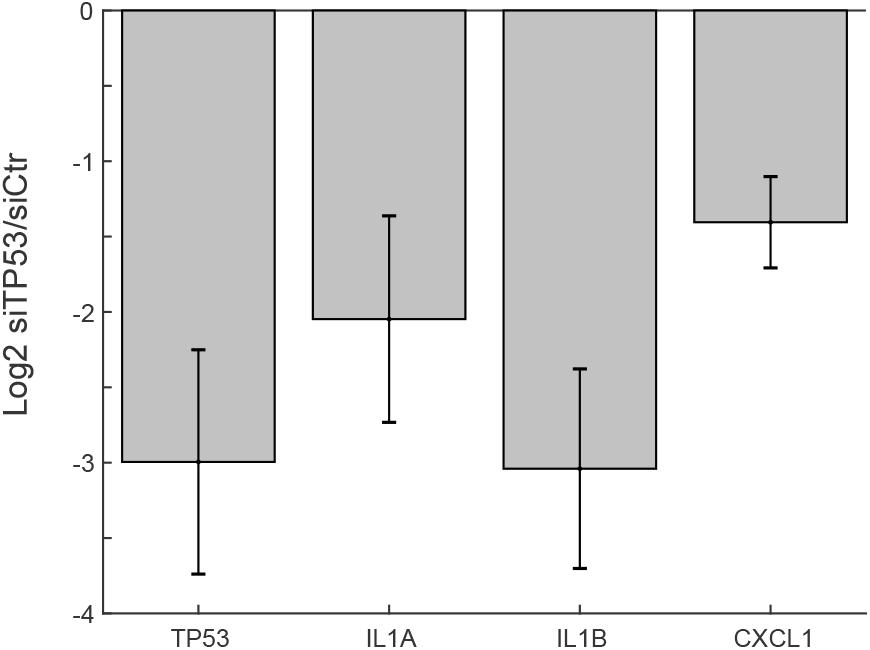
Knockdown of p53 in LOXIMVI cells reduces expression of inflammatory genes. Expression of p53, IL1A, IL1B, and CXCL1 by qPCR in cell treated with p53si compared to control siRNA (N=4).

